# Temporal shifts in microRNAs signify the inflammatory state of primary murine microglial cells

**DOI:** 10.1101/2025.04.11.648484

**Authors:** Keren Zohar, Elyad Lezmi, Fanny Reichert, Tsiona Eliyahu, Marta Weinstock, Michal Linial

**Affiliations:** Department of Biological Chemistry; Department of Genetics, Faculty of Life Sciences; Institute of Drug Research, School of Pharmacy, The Hebrew University of Jerusalem, Jerusalem, Israel

**Keywords:** Innate immune system, IL-1, RNA-seq, Purinergic receptor, inflammation, cytokines, miRBase, TarBase, CLIP-Seq

## Abstract

The primary function of microglia is to maintain brain homeostasis. However, in several neurodegenerative diseases, such as Alzheimer’s disease, the pathophysiological hallmarks that drive disease progression involve neurotoxicity and alterations in neuroinflammation. In this study, we exposed murine neonatal primary microglial cultures to external signals that mimic the in vivo stimuli caused by pathogens, injury, or toxic agents. In the presence of benzoyl ATP (bzATP) and lipopolysaccharide (LPS), we observed a coordinated increase in the expression of interleukins and chemokines. We focused on the dynamics of the differentially expressed microRNAs (miRNAs) that are statistically significant (DEMs) and tracked their post-activation dynamics. Monitoring miRNAs 3 and 8 hours (h) post-activation revealed robust changes in 33 and 57 DEMs, most of which were upregulated. The DEMs exhibiting the strongest temporal regulation included miR-155, miR-132, miR-3473e, miR-222, and miR-146b. Additionally, a strong downregulation of miR-3963 was attributed to the exposure to bzATP. Through their regulation of TNFα and NFκB signaling pathways, the identified DEMs reflect the cellular response to inflammatory signals. We incubated the activated cells with ladostigil, a neuroprotective compound that has been shown to reduce oxidative stress, inflammation, and cognitive decline. While there was no significant effect 3 h post-activation, at 8 h, a few miRNAs implicated in inflammation suppression such as miR-27a, miR-27b, and miR-23b were upregulated in a ladostigil-dependent manner. We conclude that the miRNA expression profile provides a sensitive indicator of the regulatory mechanisms underlying inflammation-related responses in microglia. We propose that primary microglia subjected to controlled activation paradigms can serve as a robust model for inflammatory states in the brain to monitor the aging brain along the progression of neurodegenerative diseases.

## Introduction

Neuroinflammation is a key factor in neurodegenerative diseases (NDDs). Diseases such as Alzheimer’s (AD), Parkinson’s (PD), multiple sclerosis (MS), and amyotrophic lateral sclerosis (ALS) are characterized by chronic inflammation, mitochondrial dysfunction, and neuronal degeneration [1, 2]. Microglia, the resident immune cells of the CNS, play a central role in responding to injury and pathological stimuli [3]. In AD, prolonged activation of microglia leads to β-amyloid clearance and synaptic loss, while the release of inflammatory mediators from microglia causes neuronal damage [4-6]. The transition of microglia from a homeostatic to an activated state in response to stressors like brain injury or pathogens [7, 8] is coupled with a morphological shift and transformation from ramified to amoeboid forms [9]. Activated microglia are categorized into pro- and anti-inflammatory states, with chronic activation leading to excessive secretion of pro-inflammatory cytokines, contributing to NDD pathology [10, 11].

Taken together, the parameters of the inflammatory response are useful markers for disease progression [12, 13]. During a chronic state, microglia secrete excessive pro-inflammatory cytokines (such as TNF-α, IL-1β, and IL-6) and reactive oxygen species (ROS). Molecular characterization of murine microglia has identified microglial-specific transcripts (∼100) that correspond to their cell identity. Among them, Tmem119, P2ry12, Siglech, and Cx3cr1 are highly expressed in resting microglia and are considered part of the homeostatic gene set. Microglial activation induces a set of molecular markers (e.g., Iba1, Cd68, Apoe, Lpl, and Cst7) together with inflammatory genes such as Ccl2, Il1b, and Nos2 [11]. Transcriptomic analysis of single-cell sequencing has expanded the list of marker genes and improved our understanding of the molecular signature underlying microglial functional states [14].

Studies on primary microglial cultures stimulated with ATP (bzATP) and lipopolysaccharides (LPS) have revealed coordinated inflammatory gene expression, including activation of the TNFα and NF-κB pathways [15]. Ladostigil, a neuroprotective compound, has been shown to reduce oxidative stress, inflammation, and cognitive decline in aging rodent brains [16]. Long-term application of ladostigil in mice restored the lowered mitochondrial potential induced in cells by H_2_O_2_ and decreased markers of oxidative stress. In microglial cultures, ladostigil inhibited nuclear translocation of EGR1 and NF-κB, phosphorylation of ERK1/2, and p38 and release of pro-inflammatory cytokines [17]. [17]. While cell lines like BV2 and S9 serve as models, primary microglia better reflect in vivo responses [18]. The microglial primary culture is amenable to pharmacological testing.

In this study, we focused on the profile of microRNAs (miRNAs) in microglial cells. MiRNAs are emerging as biomarkers for neurodegenerative diseases (NDDs) due to their role in regulating inflammation and oxidative stress, with potential applications in disease monitoring [19, 20]. However, it is not yet clear how the modulate microglial activation selectively preserve their neuroprotective functions [21]. This study examines the changes in the miRNA transcriptomic profile upon stimulation and in the presence of ladostigil. We monitor the dynamics of the activation process by focusing on the differentially expressed miRNAs and discuss these miRNAs as potential indicators of microglial inflammatory states.

## Materials and Methods

### Compounds and reagents

The cell culture reagents, including Dulbecco’s Modified Eagle Medium (DMEM), DMEM/F12, gentamycin sulfate and L-glutamine were obtained from Biological Industries (Beit-Haemek, Israel). The activation protocol included stable ATP 2ʹ-3ʹ-*O*-(4-benzoyl benzoyl) adenosine 5ʹ-triphosphate (BzATP) and lipopolysaccharide (LPS), from Escherichia coli 055:B5, purified by trichloracetic acid extraction (Sigma-Aldrich, Jerusalem, Israel). Ladostigil was a gift from Spero Biopharma (Israel).

### Preparation of microglial cultures

Primary microglia were prepared according to a previously described protocol [22]. The cells were isolated from the brains of neonatal male Balb/C mice (Harlan Sprague Dawley Inc., Jerusalem, Israel). Briefly, cells were isolated and plated for 1 week in poly-L-lysine-coated flasks. Following a dissociation protocol, non-adherent and loosely adhered cells were re-plated for 1 h on uncoated bacteriological plates, which allowed the removal of cells exhibiting slower adherence kinetics. Microglial cells were propagated and supplemented with conditioned medium containing mouse-CSF (colony-stimulating factor) and maintain in heat-inactivated fetal calf serum (FCS). For experiments, the microglial culture was washed and the medium was replaced with purified BSA for 24 h. The purity of the culture was confirmed by morphological criteria and a set of markers as previously described [23]. The cultures are estimated to be >95% microglia pure. Under such conditions, the microglial culture remains responsive for four weeks.

### Measurement of cytokines

Data presented are for microglial cells stimulated by a combination of bzATP (400 µM) and LPS (0.75 µg/mL). We showed that the concentration of LPS (0.75 µg/ml) given together with BzATP did not affect cell viability after 3 and 24 h using the MTT assay as previously described [17]. Cell lysates were tested and normalized by protein levels using BCA Protein Assay (Pierce, Meridian, Rockford, IL, USA) and cytokine ELISA assay were performed according to manufacturer’s protocols. Cells were grown to 75% confluence in 6-well plates. Measurements of cytokine secretion were made 24 h after activation in the presence of BSA (0.4 µM) using Max deluxe (Biolegend, CA, USA) commercial ELISA kits. Assays for cytokine release were conducted with 5 × 10^5^ cells per well and cytokine assays were calibrated by using internal standard curves.

### MicroRNA-seq

Microglial cultures were harvested using a cell-scraper. Total RNA was purified from ∼10^6^ cells using QIAzol Lysis Reagent RNeasy plus Universal Mini Kit (QIAGEN, GmbH, Hilden, Germany). To ensure homogenization, a QIAshredder (QIAGEN, GmbH, Hilden, Germany) mini-spin column was used. Samples were transferred to a RNeasy Mini spin column and centrifuged for 15 s at 8000× *g* at room temperature. The mixture was processed according to the manufacturer’s standard protocol. Samples with an RNA integrity number (RIN) >9, as measured by Agilent 2100 Bioanalyzer, were considered for further analysis. RNA libraries (in triplicates) were prepared according to NEBNext Small RNA Library Prep Set for Illumina (Multiplex Compatible) Library Preparation Manual. Adaptors were then ligated to the 5ʹ and 3ʹ ends of the RNA, and cDNA was prepared from the ligated RNA and amplified to prepare the sequencing library. The amplified sequences were purified on E-Gel^®^ EX 4% Agarose gels (ThermoFisher, Waltham, MA, USA, # G401004), and sequences representing RNA smaller than 200 nt were extracted from the gel. The library was sequenced using the Illumina NextSeq 500 Analyzer.

### Bioinformatic analysis and statistics

All next-generation sequencing data underwent quality control using FastQC, version 0.11.9. Adapter trimming and quality filtering were performed using Trimmomatic, version 0.39, with the AllTrueSeqPE adapter list and a minimum read length threshold of 15 nucleotides [24]. Processed reads aligned to the reference genome (GRCm38) using miRDeep2 [25]. Quantification of miRNAs was performed using miRDeep2 with miRBase v22 annotations. Trimmed mean of M-values (TMM) normalization of miRNA read counts and differential expression analysis were performed using edgeR, version 3.36.0 [26]. TMM is a between-sample method that is suitable to compare different libraries. The low variability of the TMM values within a triplicate group confirms the quality of the RNA-seq data. For differential expression (DE) analysis, genes were filtered by requiring at least three samples to have a counts-per-million (CPM) value greater than four, with an FDR q-value < 0.05. The term ‘Same’ marks changes in expression that are bounded by 33%. Partition of DEMs to clusters was done according to threshold listed in Supplementary **Table S1**.

Results of unsupervised clustering was performed using the R-base function “prcomp”. The analysis captures the maximum amount of variation in the data. Figures with added annotation were generated using the ggplot2 R package, version 3.3.5. All other statistical tests were performed using R-base functions. When appropriate, *p*-values <0.05 were calculated and considered statistically significant. Results from microglial cell experiments are presented as mean ± SD (standard deviation; see details in [15]).

Physical protein-protein interaction (PPI) set is based on STRING [27]. STRING network was used based on high PPI confidence score (>0.6). Connectivity network excluded neighborhood, gene fusion and co-occurrence as evidence. miRNet 2.0 was used for analyzing miRNA functions through network-based visual analytics. For functional analysis, the platform integrates miRNA with targets, transcription factors (TFs) and a knowledge-based graph [28]. Enrichment for miRNAs was based on miRinGO, that address indirect gene targets genes through transcription factors (TFs) according to miRNA expression in specific tissues [29]. Database of dbDEMC 3.0 (database of differentially expressed miRNAs in human cancers) covers 40 cancer types with large-scale compilation of miRNA gene expression from experiments [30].

## Results

### Release of pro-inflammatory cytokines in response of microglia to activation

Microglial function was assessed by quantifying TNF-α and IL-6 secretion after stimulation (see Methods). The baseline of secreted cytokines in untreated cells was below detection levels. TNFα and IL-6, were only detected 8 and 24 h after activation. At 8 h post activation, the absolute levels of TNF-α and IL-6 were approximately 430 and 6 pg/μg, respectively. We normalized the amount of protein released to the basal level measured for bzATP, 24 h after exposure (**Figure 1A)** The level of induction was >20 fold higher than that in the presence of bzATP. While the absolute level of IL6 was lower than that of TNF-α, the induction was approximately 50-100-fold for the combination of bzATP/LPS (**Figure 1B**). The pro-inflammatory cytokine secretion increased significantly and consistently for 48 h. However, IL-6 secretion was slightly less stable and was sensitive to cell density (at a range of 1 × 10^5^ to 5 × 10^5^ cells per well). We analyzed the transcript levels of Tnf and Il6 in untreated and the stimulated cells. Tnf was detectable at low levels in naïve cells but not Il6 in accordance with the results from secreted cytokines. While monitoring mRNA levels is valid in a few hours post activation, measurement of the secreted proteins, requires a longer time for completing ribosomal translation, folding, post modifications and trafficking. Similar to the results from **Figure 1B**, the secretion of IL-6 continues to rise, which is consistent with the increase in Il6 transcript levels. These findings indicate that the pro-inflammatory response is robust and coordinated, while the activation kinetics is unique for the different cytokines.

**Figure 1.**
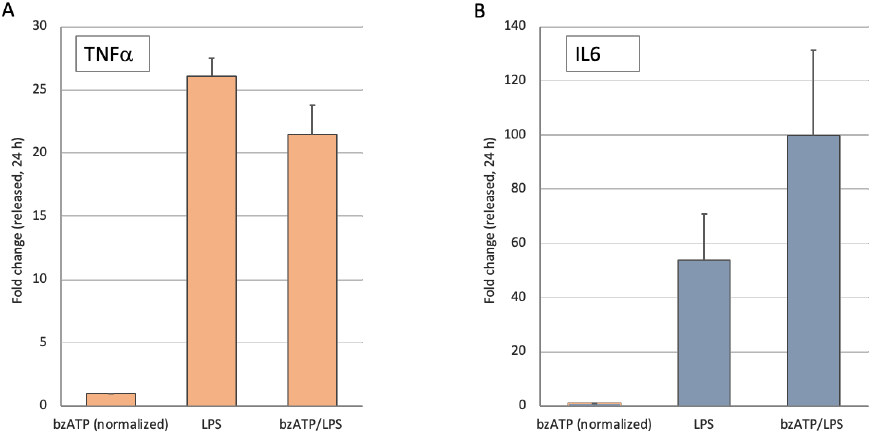
Quantitation of TNFα and IL-6 released from primary neonatal murine microglial culture following stimulation. **(A)** Relative induction of TNFα protein 24 h post stimulation. Values are normalized to the levels of TNFα in the presence of bzATP. **(B)** Relative induction of IL-6 protein 24 h post stimulation. Values are normalized to the levels of IL-6 in the presence of bzATP. Notably, the basal level of TNFα and IL-6 were below detection in unstimulated cells. Samples were collected from conditioned media supplemented with BSA, harvested, and measured 24 h after stimulation. A mean and standard deviation (s.d.) of 3 experiments in triplicates.

### The alternation in miRNA profile in the presence of bzATP is minimal

We sought to follow the temporal behavior of the activated cells by the change in the miRNA profiles. None of the 372 identified miRNAs was significantly changed 3 h after exposure to BzATP (restricted to the predetermined expression threshold, see Methods), and only a minimal change in miRNA profile was detected 8 h after exposure. Among all 372 miRNAs only three were statistically significant (i.e., DEMs), miR-146b-3p and miR-146b-5p were upregulated to a level of 1.66 and 1.41 relative to untreated cells (N.T.) while miR-3963 was downregulated. Notably, the degree of downregulation of miR-3963 was 5.8-fold in the presence of bzATP/LPS after 8 h, but did not reach the maximal level shown by bzATP alone (10.9-fold). From these results we conclude that the addition of bzATP alone was unable to activate the full response of microglial cells, but potentially primes miR-146b, and suppresses miR-3963.

### Temporal expression miRNA profiles following bzATP/LPS activation

As we observed a minimal response of pro-inflammatory cytokine secretion by exposure cells to bzATP (**Figure 1**), we analyzed the miRNA-seq results in cells that were subjected to the activation protocol by bzATP and LPS (bzATP/LPS). We tested the cells’ miRNA transcriptome at 3 h and 8 h after exposure. The experimental groups were separated using unsupervised clustering based on the miRNA expression data (Supplementary **Figure S1**). We identified 372 miRNAs that represent 345 uniquely labeled using miRNA-seq analysis. We measured the fold change relative to untreated cells and set a relaxed threshold on the fold change (log_2_FC >|0.33|). Each identified miRNA was assigned by its expression trend relative to that in the untreated cells as well as to the expression level monitored at a previous time point.

**Figure 2A** shows the partitions of miRNAs into modules according to their expression patterns. We divided all miRNAs into nine expression patterns by their paired expression at the time points (expression trends are defined as in Supplementary **Table S1)**. Each cluster is indicated by the number of miRNAs that match the expression pattern. As expected, most miRNAs (59%) were marked as unchanged even after 8 h (labeled ‘same–same’). Another 14% were characterized by a delayed response (i.e., labeled as ‘same-up’ and ‘same-down’; **Figure 2A**, black color). We found that 18% of the miRNAs were already upregulated in the short time frame (3 h post activation) with more of them displaying a transient expression wave (**Figure 2A**, orange color). Only 6 miRNAs exhibited a consistant and robust increase in expression (**Figure 2B**). These miRNAs are expected to carry the signature for maintaining the activated state of the microglia culture. The identified miRNAs include miR-146b, miR-155-5p, miR-29b-2, miR-5121 and miR-6240. For 11% of the miRNAs, expression was suppressed compared to that in naïve cells (**Figure 2A**, blue color) and the expression of miR-301a and miR-760 was monotonically downregulated (**Figure 2C**). We conclude that many miRNAs are subjected to temporal regulation during the initial hours following the activation protocol by bzATP/LPS. Expression modules for all miRNAs are listed in Supplementary **Table S2**.

**Figure 2.**
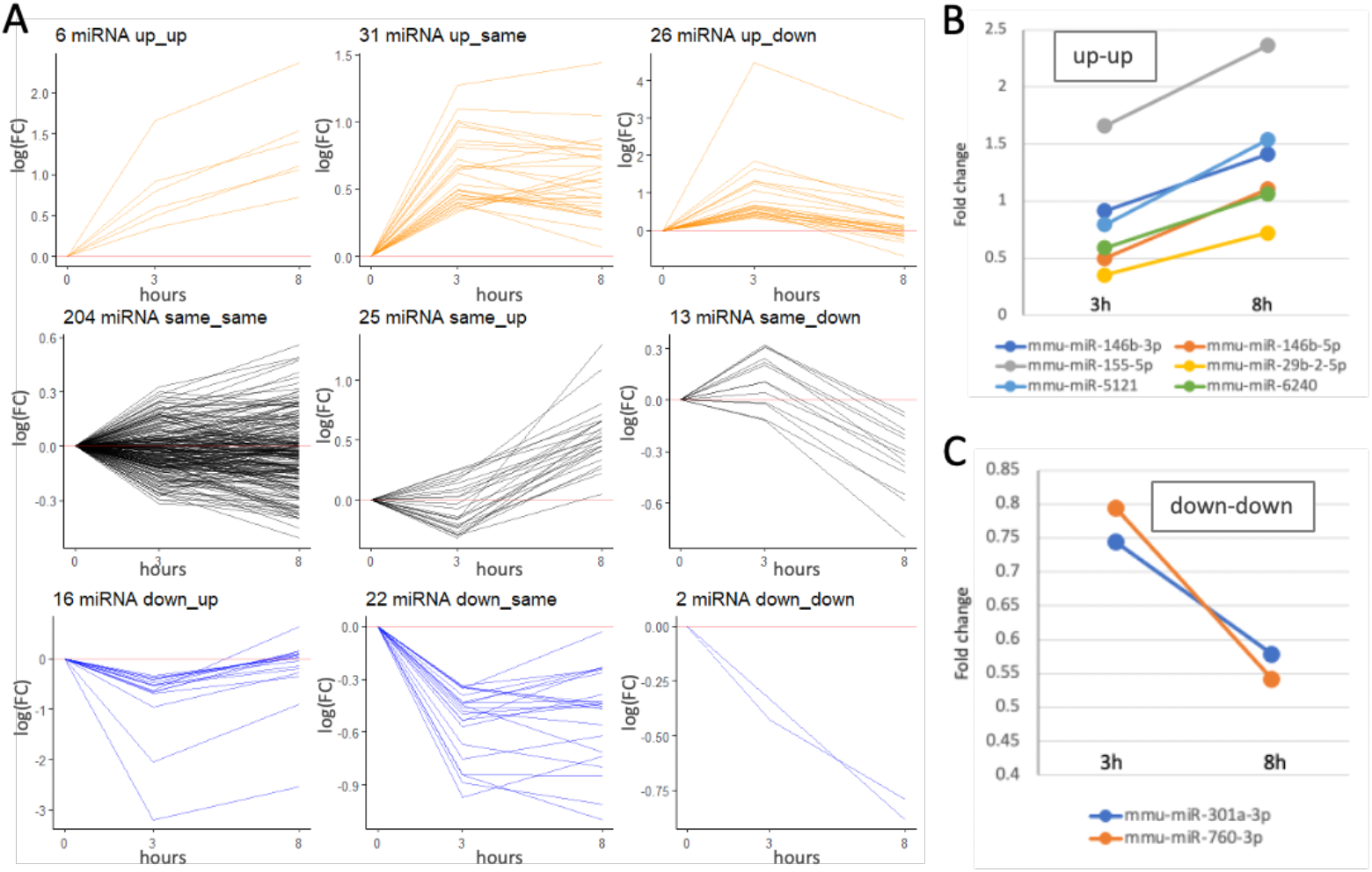
Trends in gene expression of microglial miRNAs following activation protocol with BzATP/LPS for 3 h and 8 h. **(A)** The 345 identified miRNAs are classified into nine modules based on their combined expression trend (up, down or same, and their combinations). For definition and thresholds see details in Supplementary **Table S1**). The baseline (log(FC) = 0) is shown by horizontal red line. The miRNAs associated with fast kinetics are colored for up (orange), same (black) and down (blue). The number of miRNAs that belong to each module is indicated. **(B)** The fold change of the miRNAs that are labelled ‘up-up’. **(C)** The fold change of the miRNAs that are labelled ‘down-down’. The analysis is based on the data in Supplementary **Table S2**.

### Activation of microglia by bzATP/LPS alters the miRNAs expression profile, including abundant miRNAs

Table 1 lists 20 miRNAs that are highly expressed (>100 CPM) and were significantly changed 8 h following activation by bzATP/LPS. The list shows miRNAs by their mature variants (i.e., 5p or 3p) and ranks them by the fold change relative to N.T. cells. Most of the miRNAs were upregulated, with maximal fold change seen for miR-155-5p. The expression of the most abundant miRNA, miR-21a-5p (accounts for 16.2% of all identified miRNAs), was strongly upregulated (**Table 1**, 1.59-fold). It is anticipated that even a moderate increase in expression of an abundant miRNA (albeit by ∼ 60%) may indirectly affect the stability of other miRNAs, thus impacting global cell regulation [31]. Not all abundant miRNA families were altered following activation. Although there are 20 such miRNAs that belong to the let-7 family, which accounts for >30% of all miRNAs in the cells, they remain unchanged (an exception is the downregulation of let-7b-5p, **Table 1**). We concluded that the i*n vitro* activation of the primary neonatal microglial culture is specific, resulting in substantial changes in miRNA profiles involving 15% of the abundant miRNAs (≥100 CPM). Among these differentially expressed miRNAs (DEMs) also the highly abundant miRNAs such as miR-21a, miR-146b and miR-7a.

**Table 1.** List of abundant (average CPM ≥100) DEMs 8 h post activation in the presence of bzATP/LPS

| miRNA | FDR | Fold change | Ave. CPM | Trend |
| --- | --- | --- | --- | --- |
| mmu-miR-155-5p | 2.04E-15 | 5.14 | 231.0 | Up |
| mmu-miR-146b-5p | 3.29E-16 | 2.15 | 35,262.0 | Up |
| mmu-miR-146a-5p | 3.17E-07 | 1.64 | 11,147.5 | Up |
| mmu-miR-21a-5p | 3.06E-13 | 1.59 | 161,946.4 | Up |
| mmu-miR-5099 | 8.14E-07 | 1.55 | 4,425.2 | Up |
| mmu-miR-7a-5p | 7.51E-09 | 1.50 | 16,738.0 | Up |
| mmu-miR-125a-3p | 2.87E-06 | 1.44 | 106.7 | Up |
| mmu-miR-99b-3p | 3.54E-06 | 1.40 | 523.7 | Up |
| mmu-miR-22-5p | 1.27E-04 | 1.40 | 169.7 | Up |
| mmu-miR-22-3p | 9.06E-08 | 1.39 | 3,072.8 | Up |
| mmu-miR-222-5p | 1.90E-05 | 1.37 | 243.8 | Up |
| mmu-miR-671-3p | 5.08E-03 | 1.33 | 238.8 | Up |
| mmu-miR-361-3p | 6.20E-05 | 1.33 | 330.7 | Up |
| mmu-miR-26a-5p | 2.87E-06 | 0.79 | 15,464.0 | Down |
| mmu-miR-101a-3p | 2.07E-05 | 0.79 | 825.0 | Down |
| mmu-miR-504-5p | 9.77E-03 | 0.78 | 114.8 | Down |
| mmu-miR-9-5p | 2.87E-06 | 0.77 | 2,290.4 | Down |
| mmu-miR-378a-3p | 1.47E-07 | 0.76 | 4,776.9 | Down |
| mmu-let-7b-5p | 1.95E-03 | 0.76 | 4,927.1 | Down |
| mmu-miR-3968 | 7.01E-03 | 0.73 | 142.6 | Down |

Applying the same analysis for the miRNAs at an earlier phase of activation (3 h) showed that only 10 upregulated miRNAs were identified and none were downregulated. Among these DEMs we identified miR-21a and miR-146b, as well as abundant miRNAs that are only significant at the early phase of the activation (e.g., miR-125a, miR-125b; Supplementary **Table S3)**. We conclude that the establishment of fully activated microglia is reflected by the temporal specificity in miRNA profiles.

### Dynamics of differentially expressed miRNAs (DEMs) by bzATP/LPS activation

Activation of the primary microglial culture by bzATP/LPS changed the expression profile of miRNAs. Of the 372 mapped miRNAs that were expressed in substantial amounts, 38.8% are expressed with ≥100 CPM (counts per million). Mapping led to minor duplication (with 345 uniquely labeled mature miRNAs defined also by their 5p and 3p arms from the precursor pre-miRNAs. **Figure 3A** shows that the majority of miRNAs have a low expression, with 50% of the miRNAs below 50 CPM and only 14 with >20,000 CPM. This set accounts for more than 73% of all cellular miRNAs (Supplementary **Table S3**. We report that as in other cells, the miRNA expression level in microglial cells ranges over 5 orders of magnitude.

**Figure 3.**
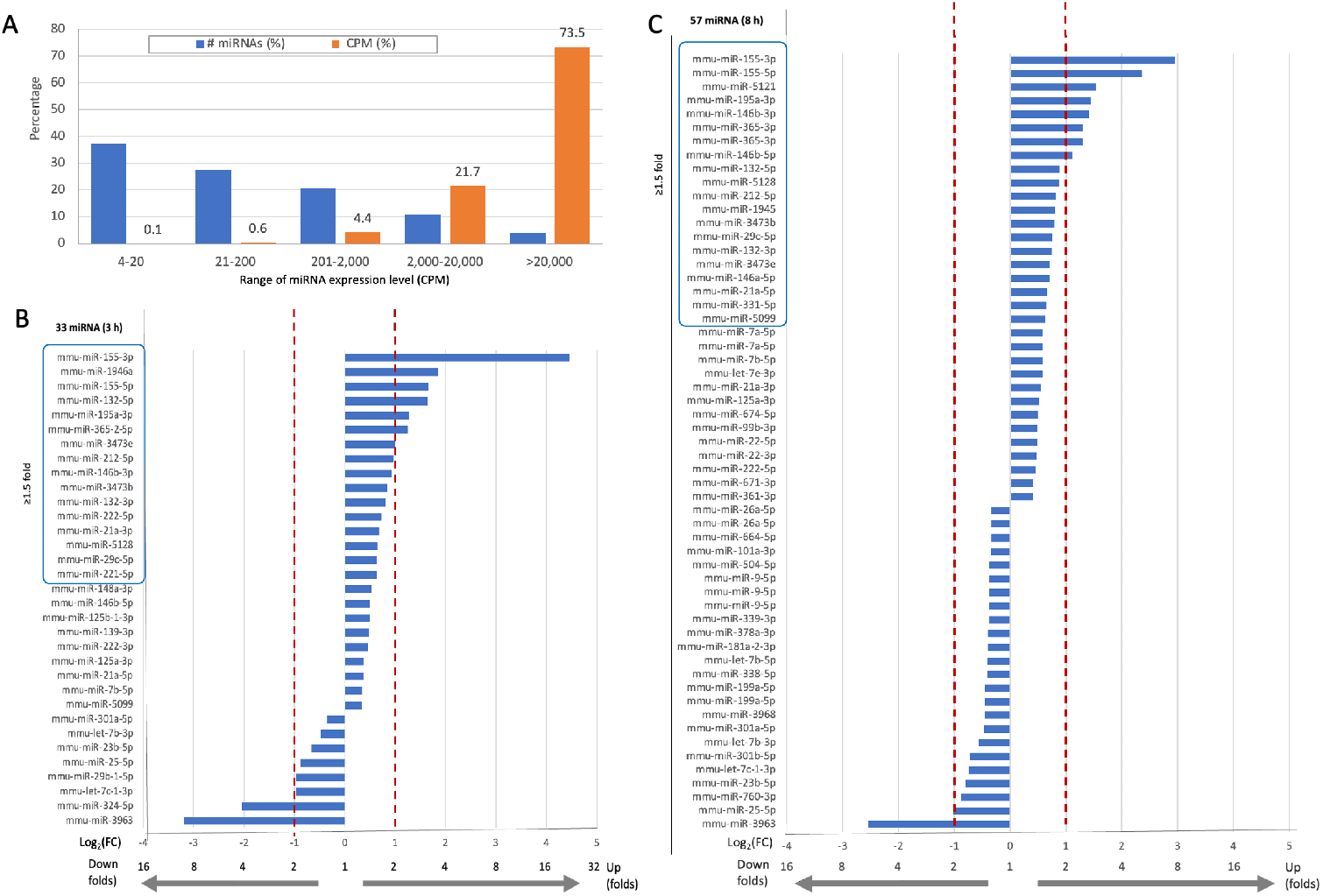
A global view of miRNAs statistics in microglia culture and time-dependent DEMs following activation. **(A)** Partition of the 372 identified miRNAs by their number of miRNAs by their expression level bins (orange, CPM) and total counts (blue). **(B)** A list of 33 significant DEMs 3 h post activation. **(C)** A list of 57 significant DEMs 8 h post-activation. The lists are sorted by fold change values. Vertical red dashed lines mark the differential expression by 2-fold, for underregulated and upregulated expression trend. The red frame around the miRNA names marks the miRNAs that were differentially expressed by ≥1.5-fold.

In all cells, the post-transcriptional regulation mediated by miRNAs on translational arrest is expected to be fast and transcriptionally independent [32]. We therefore tested the dynamics of miRNA at two time points following stimulation by bzATP/LPS (**Figure 3**). Normalized expression levels of all identified miRNAs are listed in Supplementary **Table S4**. There were 33 and 57 DEMs among the 372 miRNAs that met the statistically significant threshold (FDR ≤0.05) for the short-term (3 hours, **Figure 3B**) and long-term (8 hours, **Figure 3C**) activation protocols, respectively, while 50% of them had a fold change of ≥|1.5|.

### A small set of temporally responsive miRNAs dictates the establishment of the fully activated microglia

Inspection of the results from the miRNAs that are consistently upregulated according to the set of time-dependent unique DEMs allowed us to infer the contribution of miRNAs in establishing the activated microglial state, but also highlighted examples of transient expression. **Figure 4** shows the DEMs along the activation timeline. **Figure 4A** present a Venn diagram with most DEMs identified for the 3 h and 8h activation paradigms (Supplementary **Figure S2**). Only 9 miRNAs are unique to the early time point. Furthermore, more than half of the DEMs at 8 h are also unique (27 DEMs), with similar numbers of up-and downregulation. **Figure 4B** (left) illustrates instances of DEMs where their fold change was maximal at the early time point. For example, the miR-155-3p was upregulated by 22-fold at 3h and 7.6-fold at 8 h of activation. An opposite trend where the fold change at a later time point continues to increase, is shown (**Figure 4B**, right). The miR-132-5p and miR-155-3p show maximal expression at 3 h and remain substantially high also at 8 h, while that of miR-155-5p and mir-146b are maximal 8 h post activation. These miRNAs exhibit strong temporal sensitivity (≤30% difference between the two time points. We concluded that these miRNAs are of a special interest as their levels may dictate a time-dependent regulation of the microglial culture that was shifted to new inflammatory states.

**Figure 4.**
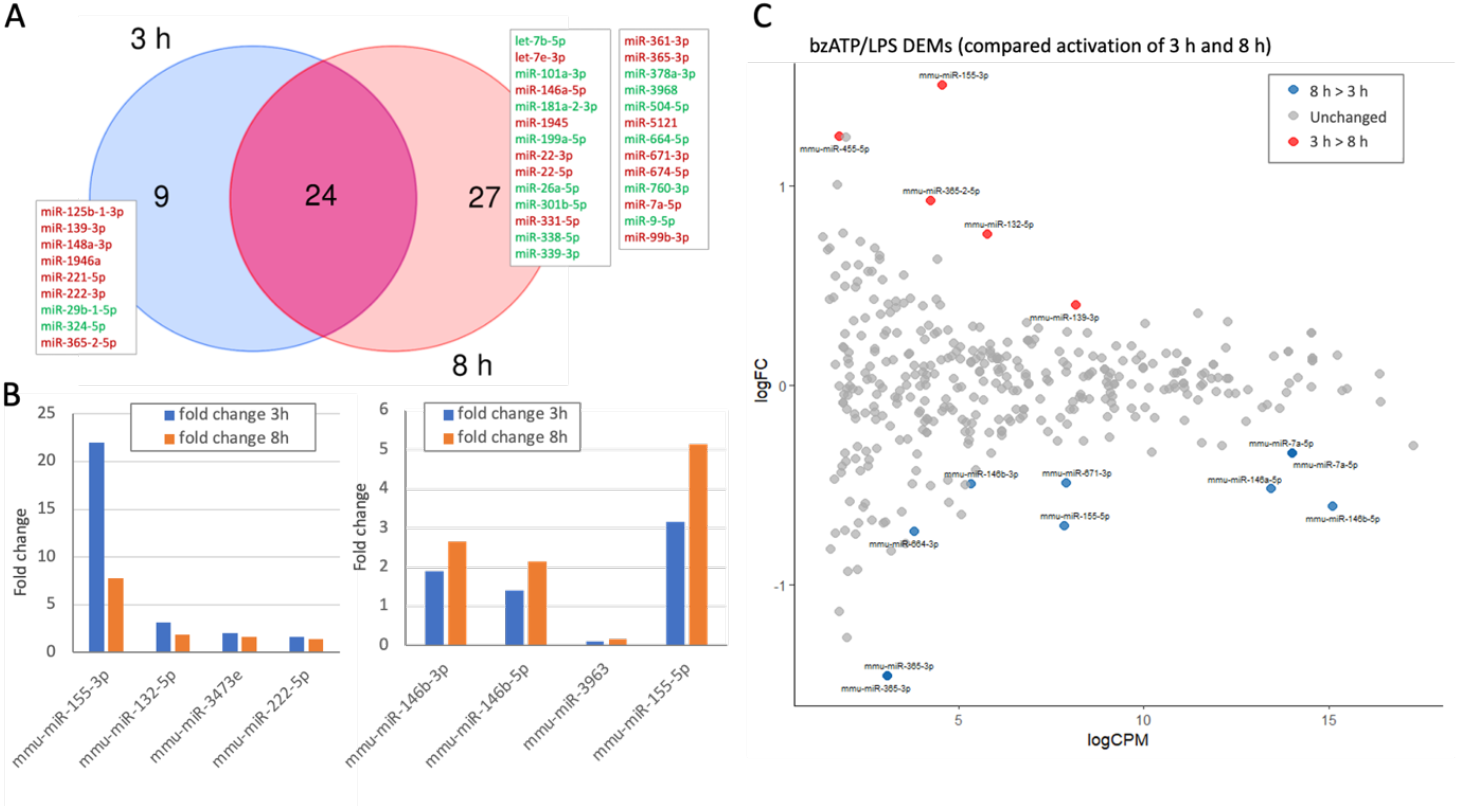
Dynamics of DEMs following activation of microglial culture. **(A)** Venn diagram of the DEMs of two time points (3 h and 8 h). The unique set that are not shared by both time points are listed. Included all significant DEMs (FDR ≤0.05) and log10(FC) of >|0.33| relative to N.T. cells. The listed names of DEGs are colored as up-and downregulated DEMs by red and green font, respectively. **(B)** Based on the overlap in A, representative DEMs are listed with a change difference >30% between the 3 h and 8 h of cell activation relative to N. T. The listed DEMs show ‘up-down’ (left) and ‘up-up’ (right). **(C)** MA plot of the log2(FC) relative to the CPM (in log) for the 372 identified miRNAs. The temporal DEMs (T-DEMs) are colored by the change expression trends with expression of 8 h > 3 h in blue and 3 h > 8 h in red. Gray are miRNAs that showed no temporal expression pattern. The change in expression was determined by a threshold of log2(FC) > |0.33| and an FDR q-value < 0.05.

To further test the dynamics of miRNAs, we focused on miRNAs that were significantly altered between the two time points during microglia activation (3 h and 8 h). **Figure 4C** displays an MA plot of time-dependent fold change versus mean expression post activation (Supplementary **Table S5**). We identified 15 miRNAs (4%) that are temporal DEMs (T-DEMs), with 5 that are maximally changed at the 3 h time point (marked red) and 10 that were further altered at a later time point (8 h, colored blue). The direct time-dependent comparison highlighted several miRNAs that are more sensitive to the dynamics of the activation process. This includes miR-455-5p, mir-365-2-5p, and miR-139-3p. An opposite trend (i.e., maximal expression occurs at 8 h post activation) included the following T-DEMs: mir-146b-3p, miR-664-3p, miR-671-3p, miR-155-5p, miR-7a-5p, miR-146a-5p and miR-365-3p. We observed that miR-365 showed a temporal dynamic, while different variants of miR-365 exhibited an opposite temporal trend (miR-365-2-5p and miR-365-p).

### Temporal expression of a set of miRNAs is coupled with differentially expressed inflammatory genes

The exposure of the cells to bzATP/LPS led to a group of miRNAs to change their expression (**Figure 4C**). We tested whether these T-DEMs affected the establishment of the microglial inflammatory state. We mapped miRNAs to their appropriate targets and limited our analysis to miRNA-mRNA pairs that were experimentally validated. The list of T-DEMs (Supplementary **Table S3**) includes miRNAs that are classified as early and late responders, based on the time point of their maximal expression (3 h or 8 h post-activation). We merged T-DEMs with RNA-seq data that were collected from microglia under identical activation conditions [15] and used miRNet 2.0 platform by its multiple-modality capability. The list of mRNA-seq results included 7,970 genes (FDR ≤0.05, filtered for coding genes) among which 149 were identified as direct targets in microglia (Supplementary **Table S6**). The rest of the genes were either not expressed in microglia or failed our thresholds. We focused on the 25 genes with a substantial temporal expression change (T-DEGs with log2(FC)<|1|). All listed miRNA-target pairs shown were validated experimentally, and were identified of microglia as T-DEGs (**Table 2**).

**Table 2.**
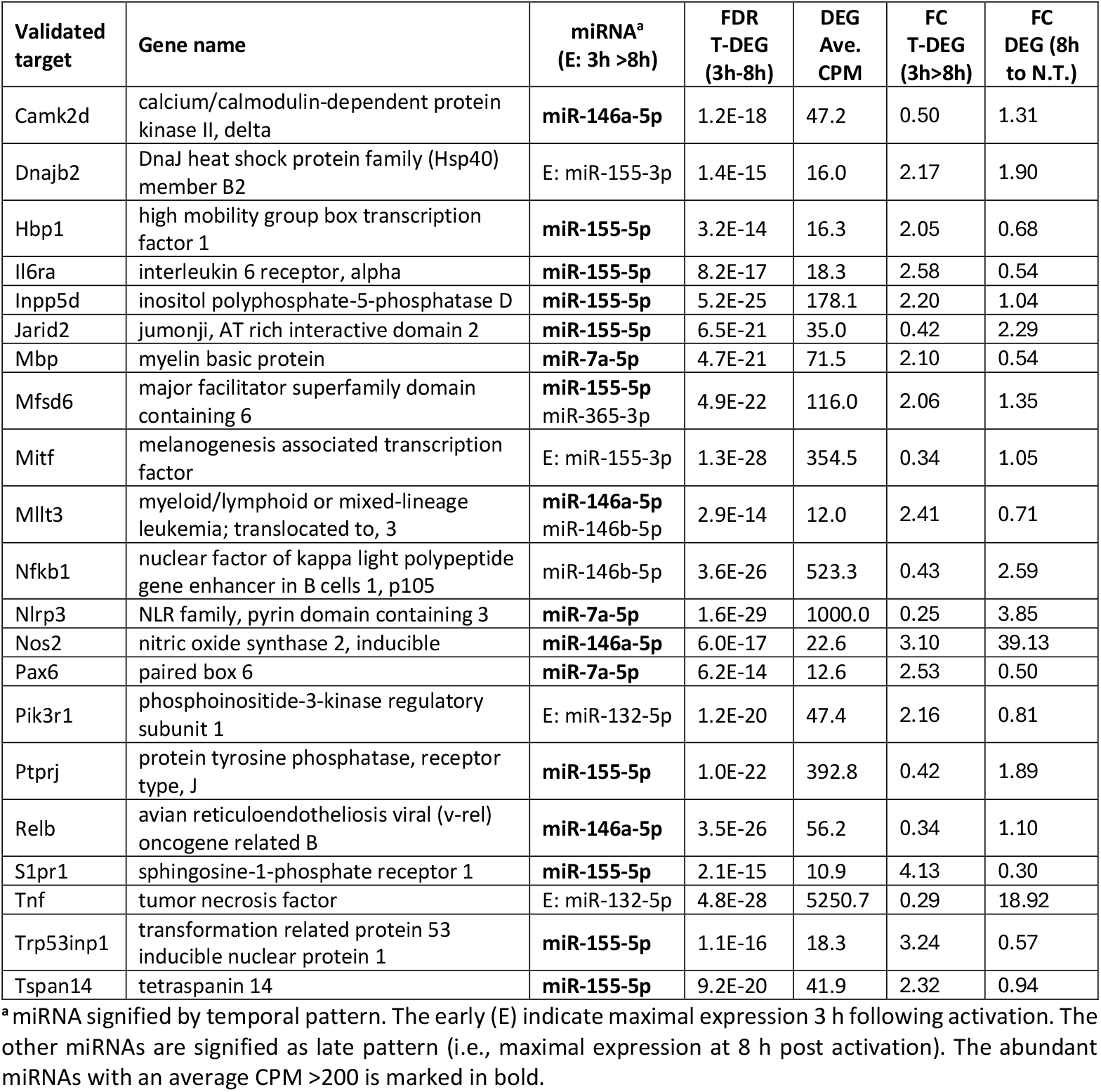
T-DEMs and their experimental validated targets based on bzATP/LPS microglia RNA-seq data.

**Table 2** lists 21 of these miRNA-targets (filtered by an expression threshold of ≥10 CPM) along with their statistical properties. Several observations emerged regarding the potential impact of miRNAs on the direct targets: (i) Most listed miRNAs are abundant (>200 CPM, bold). Extremely abundant miRNAs include miR-7a-5p and miR-146a-5p with expressed levels of 16,738 and 11,147 CPM, respectively. (ii) miR-155 is associated with many of the targets (11 of 21). (iii) Some of the target genes are very highly expressed in microglia. Examples include Tnf (CPM 5250.7), Nlrp3 (CPM 1000) and Nfkb1 (CPM 533.3). (iv) Among the listed targets, five were strongly upregulated compared to naïve cells (DEMs with log2(FC)<|1|), with Nos2 and Tnfa showing fold change of 29.13 and 18.92, respectively.

**Figure 5A** shows the results of miRNet 2.0, centered around a small set of T-DEMs. It is evident that miR-155-5p is a major hub, with a large number of paired mRNA genes, and some genes are regulated by more than one miRNA. Revisiting the results from miR-155-5p shows that it is strongly upregulated (22-fold) just 3 h of post-activation and remains high at 8 h. The direct targets of miR-155 are Socs1 (suppressor of cytokine signaling 1) and Inpp5d (inositol polyphosphate-5-phosphatase D), leading to enhanced NF-ΚB signaling and inflammation. In addition, Tnf acts as a hub in the inflammatory network (**Figure 5B**). It is paired with T-DEM miR-132-5p which is maximally expressed 3 h after exposing the cells to bzATP/LPS (**Table 2**). The expression at 8 h of the Tnf transcript is 3.4-fold higher than at 3 h. The miR-132 is highly conserved between humans and mice and has been shown as a regulator of neural signaling, where its dysregulation contributes to axonal damage. A direct interaction of miR-132 with Tnf transcript remains questionable. Still, miR-132-5p negatively regulates the release of TNF, possibly by targeting upstream regulators like NF-κB [33]. Although speculative, it is likely that the presence of miR-132-5p at early time point, and its reduced amounts at later stages, led to a delayed accumulation of Tnf transcripts in activated microglial cultures. We show that miR-146a-5p is paired with the Nos2 (nitric oxide synthase 2, **Table 2**). Overexpression of miR-146a-5p in the BV2 microglial cell line reduced the expression of iNOS, encoded by Nos2 gene [34]. A similar effect was observed in miR-146a knockout mice, which resulted in the overproduction of pro-inflammatory cytokines (e.g., IL-1β, TNFα, IL-6) [35]. These results confirm that miR-146a-5p acts to attenuate the microglial inflammatory state.

**Figure 5.**
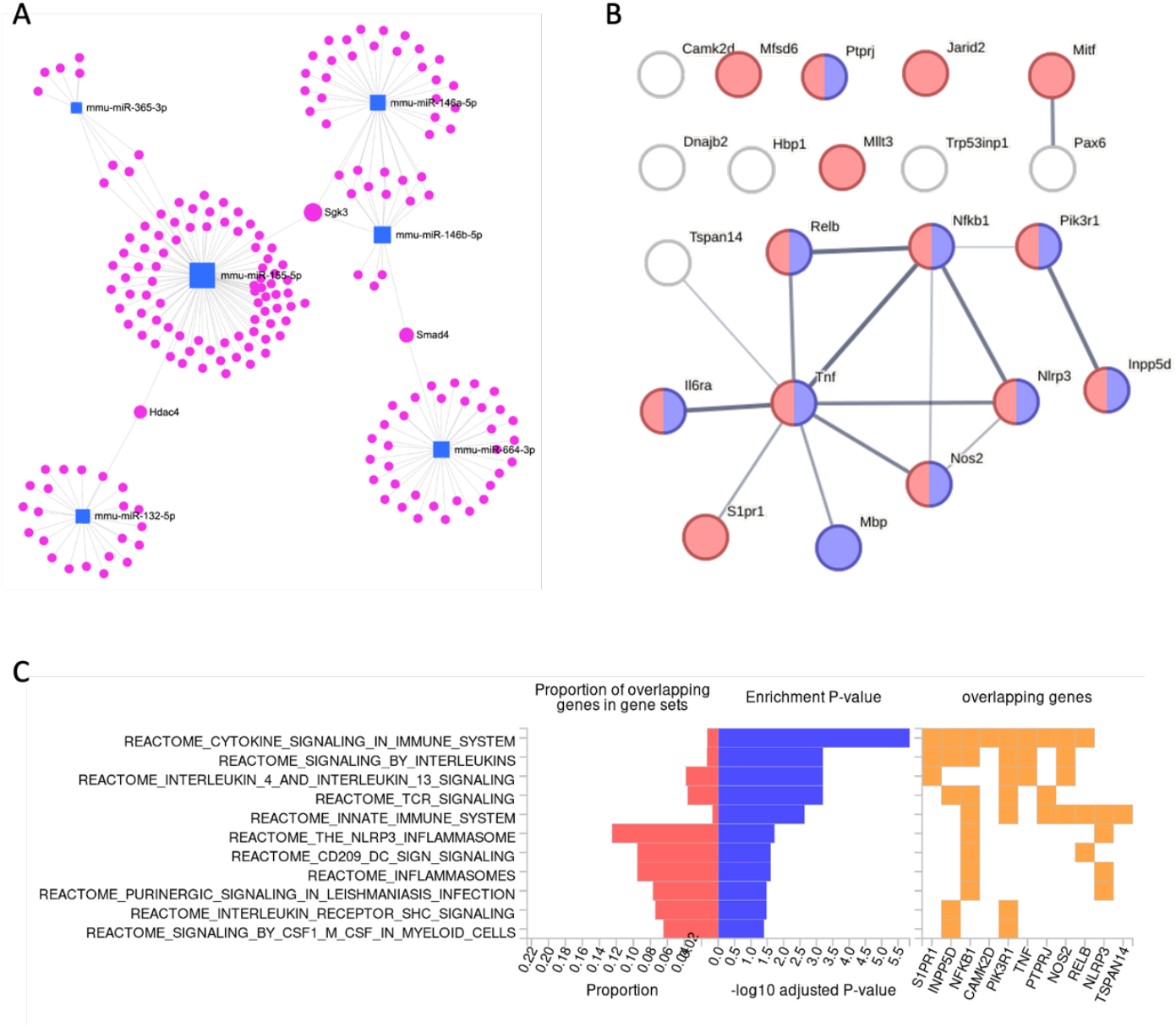
Network view for T-DEMs and gene targets. **(A)** miRNet 2.0 view for the 6 identified T-DEMs (blue square) and their experimentally validated targets (pink marker, 244 genes). Genes connecting several miRNAs are labelled. The dominant connection of miR-155-5p is evident. **(B)** STRING view for a set of 21 protein coding targets (T-DEGs, **Table 2**). The protein-protein interaction (PPI) network is significant (PPI enrichment p-value: 0.00986). Enrichment of gene ontology biological process (GO_BP) of Immune system process (GO:0002376) and Regulation of cytokine production (GO:0001817) are colored red and blue, respectively (p-value 3.4 e-06). **(C)** The functional enrichment of the listed genes (21 genes, Table 2) by Reactome pathways. From left to right: The fraction of gene input (red); Significant pathways by Reactome are sorted by p-values (blue); The overlapping genes for each of Reactome’s enriched pathways (orange).

Nlrp3 is one of the genes most strongly expressed after microglial activation. The assembly of the NLRP3 inflammasome leads to the release of the pro-inflammatory cytokine, IL-1β. While no direct effect of miR-7a-5p on Nlrp3 was shown in microglial cells. miR-7a-5p is highly expressed and known to play a role in modulating neuroinflammation by directly targeting Nlrp3, resulting in reduced pro-inflammatory cytokine production. Over half of all targeted genes to T-DEMs are annotated as ‘regulation of cytokine production’ (GO:0001817; enrichment p-value 3.4e-06). We conclude that miR-146a-5p acts to attenuate the microglial inflammatory state. While **Figure 5A** presents overall interactions of miRNAs with validated target genes from any tissue, **Figure 5B** shows protein-protein interaction (PPI) network of target T-DEGs, as identified by RNA-seq from bzATP/LPS activated microglia [15]. Many of the listed genes belong to cytokine release and immune-related categories. We further tested the miRNA-target pairs (**Table 2**) by examining the pathways represented in Reactome [36]. **Figure 5C** presented an enrichment analysis for the set of pathways that dominate the input genes. We show that cytokine signaling, interleukins, and TCR pathways are very significant. Other pathways indicate that a large fraction of the input gene list was implicated in the NLR3 inflammasome.

### Ladostigil induced miRNAs that may serve as mediators in suppressing inflammation

Analysis of miRNA profiles in the activation protocol Indicated that the system is suitable for pharmacological manipulation and analysis [37]. Accordingly, we examined the effect of ladostigil on miRNA profiles. Ladostigil, an aminoindan derivative [38] has been shown to reduce the production and secretion of pro-inflammatory cytokines [17]. RNA-seq analysis revealed that Egr1, Egr2 (Early growth response protein 1 and 2) and several metalloproteinases (MMPs) were upregulated following microglia activation protocol while incubation of ladostigil significantly reversed this trend [17, 37]. We wanted to determine whether miRNAs could be the mediators controlling the overall reduction in the inflammatory state of microglia. miRNAs may exert their function by several modes of action including cellular relocation [39], loading into AGO proteins for stabilization, indirect competition with other RNAs [40]. In a a short time-window, the miRNA profile was compared between to that in N.T. (with 2 h incubation with ladostigil) and 3 h following full activation. We found that none of the 372 identified miRNAs was significantly altered (**Figure 6A**). We concluded that miRNA expression changes were not involved in the immediate response to ladostigil. However, at 8 h ladostigil, significantly upregulated 4 miRNAs: miR-23b-5p (1.48-fold), miR-27a-5p, miR-27b-5p (1.27-1.28-fold; **Figure 6B**, Supplementary **Table S7**) and miR-365-2-5p (1.93-fold). While the role of miR-365-2-5p is not known, the other miRNAs have previously been implicated in inflammation suppression (**Table 3**). Despite the relatively moderate fold changes in miR-27a and miR-27b, they are highly abundant and thus may be involved is suppression of multiple targets (average amounts in microglia is 532.2 and 167.5 CPM, respectively). Specifically, miR-27b counteracted the effects of TNFα and was linked to restoring mitochondrial function, reducing apoptosis, and the Akt/Foxo1 pathway [41]. Others have shown that changes in miR-23b are associated with PD [42], and have been implicated in the oxidative stress response [43] and in suppressing α-synuclein expression. In mice with induced sepsis, miR-23b-5p was reduced, coupled with the elevation in ADAM10 and other MMPs. Expression of metalloproteases increased in the activated microglia, was suppressed by ladostigil, together with the decrease in the levels of inflammatory cytokine (TNF-α, IL-1β, IL-6) [17].

**Figure 6.**
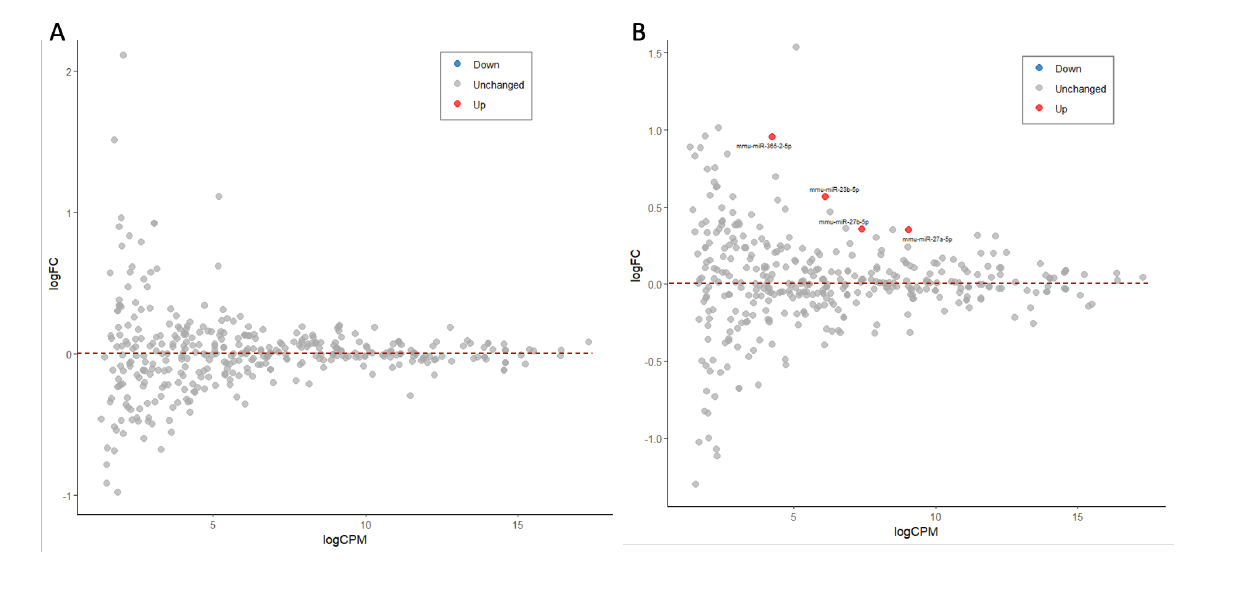
The effect of ladogtigil on miRNA expression. MA plots of the log2(FC) relative to the CPM (in log) for the 372 identified miRNAs in a fully activated setting (bzATP/LPS). **(A)** 3 h post activation relative to N.T. **(B)** 8 h post activation relative to N.T. The miRNAs that were upregulated by ladostigil are marked in red font. Gray color are miRNAs that showed no significant expression change. The change in expression was determined by a threshold of log2(FC) > |0.33| and an FDR q-value <0.05. The dashed lines mark stable expression.

**Table 3.**
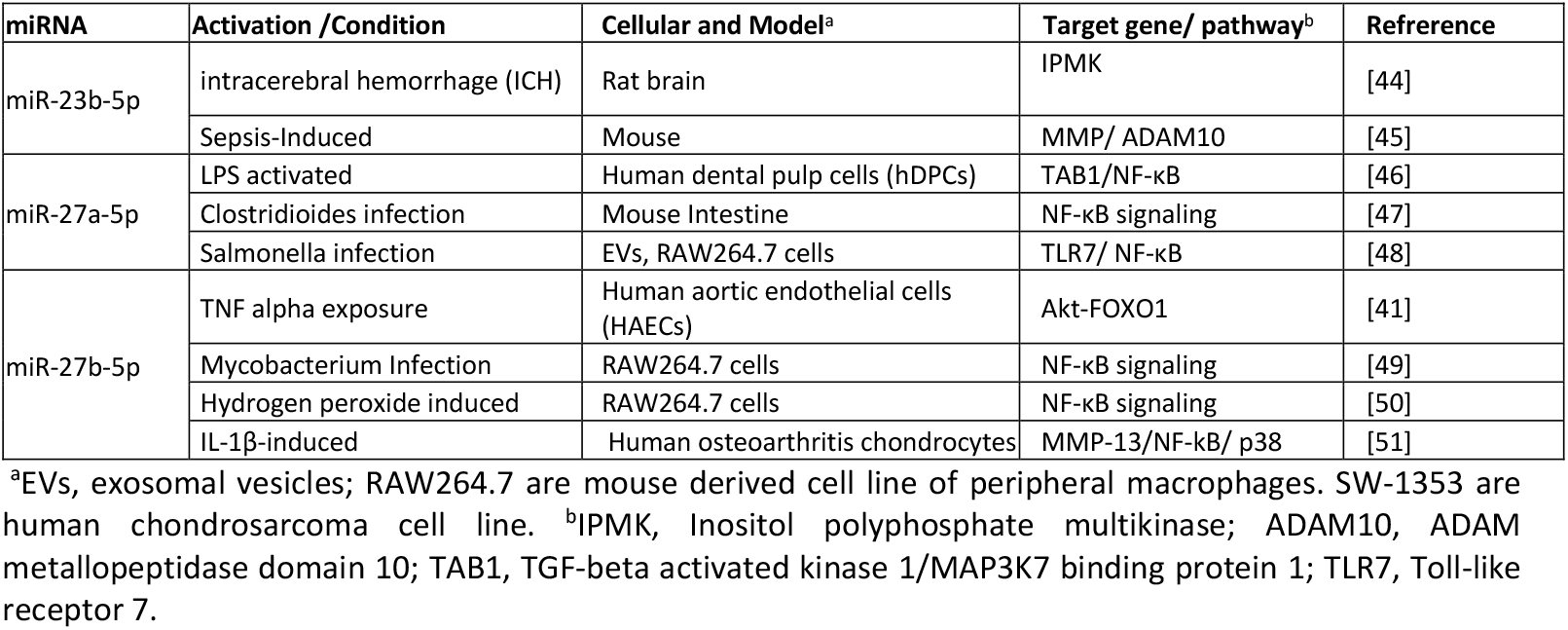
Upregulation by ladostigil of miRNAs that act in reducing inflammation in multiple model systems

| miRNA | Activation /Condition | Cellular and Model<sup>a</sup> | Target gene/ pathway<sup>b</sup> | Reference |
| --- | --- | --- | --- | --- |
| miR-23b-5p | intracerebral hemorrhage (ICH) | Rat brain | IPMK | [44] |
|  | Sepsis-Induced | Mouse | MMP/ ADAM10 | [45] |
| miR-27a-5p | LPS activated | Human dental pulp cells (hDPCs) | TAB1/NF-κB | [46] |
| | Clostridioides infection | Mouse Intestine | NF-κB signaling | [47] |
| | Salmonella infection | EVs, RAW264.7 cells | TLR7/ NF-κB | [48] |
| miR-27b-5p | TNF alpha exposure | Human aortic endothelial cells (HAECs) | Akt-FOXO1 | [41] |
| | Mycobacterium Infection | RAW264.7 cells | NF-κB signaling | [49] |
| | Hydrogen peroxide induced | RAW264.7 cells | NF-κB signaling | [50] |
| | IL-1β-induced | Human osteoarthritis chondrocytes | MMP-13/NF-κB/ p38 | [51] |

<sup>a</sup>EVs, exosomal vesicles; RAW264.7 are mouse derived cell line of peripheral macrophages. SW-1353 are human chondrosarcoma cell line. <sup>b</sup>IPMK, Inositol polyphosphate multikinase; ADAM10, ADAM metallopeptidase domain 10; TAB1, TGF-beta activated kinase 1/MAP3K7 binding protein 1; TLR7, Toll-like receptor 7.

**Table 3** summarizes the current knowledge on the link between the overexpression of miRNAs and their potential targets that drive inflammation reduction. We conclude that ladostigil induces a restricted set of miRNAs that enhance the ability of the cells to cope with oxidative stress and the induction of pro-inflammatory signature of the treated cells. The central role of miRNAs in attenuating NF-kB signaling (**Table 3**) is in accord with the significance of this pathway in the modulation by ladostigil in other cellular systems (e.g., [37]).

## Discussion

Our study focused on changes in miRNAs in primary neonatal purified cultures of microglia that exhibit a strong response to external signals such as bzATP and LPS. These stimuli mimic the microglial microenvironment upon exposure to pathogens causing substantial cell death. The contribution of miRNAs to the inflammatory environment has been extensively studied in neurodegenerative diseases (NDDs). Dysregulation of miRNAs in microglia has been reported, in the ALS mouse model (with mutated SOD1), along with their response to inflammatory signals, including microglial NLRP3 inflammasome activation [52]. miR-365 and miR-125b were among the miRNAs significantly upregulated in ALS microglia, [53]. The study showed that miR-365 interferes with the interleukin-6 (IL-6) pathway, while miR-125b affects the STAT3 signaling pathway. These interactions led to increased production of TNFα, contributing to neurodegeneration in ALS. Similar to our findings, in the model of ALS, miR-22, miR-155, miR-125b, and miR-146b were upregulated [53].

Our results are consistent with the centrality of miR-155 in the inflammatory phenotype of M1 macrophages [54, 55]. Using real-time PCR, it was confirmed that miR-155 contributes to the suppression of mRNA targets that lead to the induction of iNos, IL-1β, Tnfα, IL-6, and IL-12. Among the direct targets of miR-155 are Inpp5d, Ptprj, and other transcripts that were also identified in our study as T-DEGs (**Table 2**). The contribution of chronic neuroinflammation to major NDDs, including AD, (PD), ALS, and multiple sclerosis (MS), is reflected by the miRNA signature from microglia [42]. We suggest that temporal analysis of cellular miRNAs can be utilized as a sensitive indicator for NDD progression and also to develop new therapeutic strategies for modulating the neuroinflammatory state.

Chronic treatment with ladostigil in aging rats attenuated some aspects of neuroinflammation, including the upregulation of the Adora 2 ATP receptor together with memory decline [9]. In microglial cultures, activation by bzATP /LPS led to the upregulation of Egr1, Egr2, and PDGF-β, which are linked to P2×7R signaling and neuroinflammation [17]. The upregulation of transcripts such as Egr1 by ladostigil highlighted its role in suppressing oxidative stress and MAPK pathways. We suggest that the observed miRNA signature supports the importance of ladostigil in counteracting neuroinflammatory processes, most likely through the suppression of oxidative damage [37]. Such regulation by ladostigil was validated in the aging rat brain and in the prevention of memory decline. We show that the specific miRNAs that were induced by ladostigil were shown in model organisms and cellular systems to suppress inflammation, most likely by the activation the NF-kB signaling (**Table 3**).

An apparent limitation of our cellular system is the lack of any crosstalk between the microglial culture and other cell types. Cell communication is expected to occur between microglia, neurons, and other cells in the central nervous system (CNS) (e.g., astrocytes, endothelial cells, stem cells). Exosomes and additional types of extracellular vesicles (EVs) serve as an added layer of cell communication [56]. Many of the miRNAs reported in this study were identified as circular miRNAs (e.g., in serum, plasma, CSF) across a wide range of CNS disorders [57]. For example, in mouse models, astrocyte-derived exosomes were shown to regulate microglial responses following traumatic brain injury (TBI). Isolated exosomes carry miR-873a-5p, which inhibited ERK and NF-κB signaling in microglia, thus reducing neuroinflammation and accelerating the repair process [58]. Increased levels of EV-associated miRNAs miR-21, miR-146, miR-7a, and miR-7b of were found in injured mouse brains [59]. It was suggested that the source of miR-21 is neurons near the lesion site, and consequently, EVs altered the response of microglia. In our system, a similar set of miRNAs was upregulated by bzATP/LPS without any trigger from neurons or other cell types. We suggest that even a moderate increase in the expression levels of miR-21, miR-146a, miR-146b, and miR-7a can cause a shift in cellular miRNA regulation due to their extreme abundance (**Table 1**). Changes in the relative abundance of miRNAs may activate an indirect effect on the availability of many other less abundant miRNAs [31].

Exosome-based communication between resting and activated microglia was validated in a mouse model, of retinal angiogenesis. It was confirmed that miR-155-5p within the exosomes caused activation of the NF-κB pathway [60]. In another system, miR-181a-3p from mesenchymal stem cells (MSCs) affected oxidative stress in PD. Using SH-SY5Y neuroblast-like cells as a model for drug-induced PD, it was shown that miR-181a-3p was transferred via EVs from MSCs to the SH-SY5Y cells, where it affected the p38 MAPK pathway by inhibiting EGR1. We argue that the attenuation of oxidative stress underpins the mechanism of action of ladostigil [37], which can be manifested directly or through EV-mediated miRNA cellular communication. The regulatory impact of miRNAs within EVs on intercellular communication is strongly dependent on amounts and targets’ stoichiometry [61]. Currently, circular miRNAs and the molecular content of exosomes are considered attractive biomarkers. The temporal expression of miRNAs in cells are promising non-invasive indicators of cellular inflammatory states. Our findings provide insights into miRNA-mRNA regulatory networks in regulating the inflammatory state of primary microglia, offering potential therapeutic targets. Dysregulation of miRNAs, especially the most abundant ones, suggests they can serve as early biomarkers for oxidative stress-induced neuronal damage in NDDs, CNS disorders, and other brain pathologies.

## Abbreviations

AD: Alzheimer’s disease
ATP: Adenosine triphosphate
BSA: Bovine serum albumin
BzATP: 2’-3’-O-(4-benzoyl benzoyl) adenosine 5’-triphosphate
DEG: Differentially expressed genes
DEM: Differentially expressed miRNAs
DMEM: Dulbecco’s modified Eagle medium
FC: Fold change
FDR: False discovery rate
GO: Gene ontology
h: Hours
LPS: Lipopolysaccharide
MAPK: Mitogen-activated protein kinase
NF-κB: Nuclear factor kappa-light-chain-enhancer of activated B cells
N.T.: Not treated
RNAseq: RNA sequencing
TMM: Trimmed Mean of M-values normalization of RNA

## Acknowledgement

We thank the members from the Linial’s lab for useful discussion. We thank Slomo Rotshenker for his support in developing the microglial culture protocol. We thank the Clore Scholars Program for the fellowship and support to K.Z.

## Data availability

RNA-seq data files for miRNAs were deposited in ArrayExpress under the accession E-MTAB-14921.

