## Supplementary material for "Temporal shifts in microRNAs signify the inflammatory state of primary murine microglial cells": Fig. S1, Fig. S2

**List of supplementary information**

Supplementary **Figure S1.** Dendogram of miRNA-seq by experimental groups

Supplementary **Figure S2.** MA scatter plots for 3 h and 8 h relative to N.T.

Supplementary **Table S1.** Thresholds used for DEMs

Supplementary **Table S2.** Expression modules for all miRNA

Supplementary **Table S3.** DEM for LPS relative to N.T. at 3 and 8 hours post activation

Supplementary **Table S4.** The normalized amounts of miRNA by TMM

Supplementary **Table S5.** DEM comparing LPS 3 h and 8 h.

Supplementary **Table S6.** List of miRNet2.0 mapping for 11 T-DEMs

Supplementary **Table S7.** DEM for LPS 8h with ladostigil relative to LPS 8h.

**Supplementary Figures**

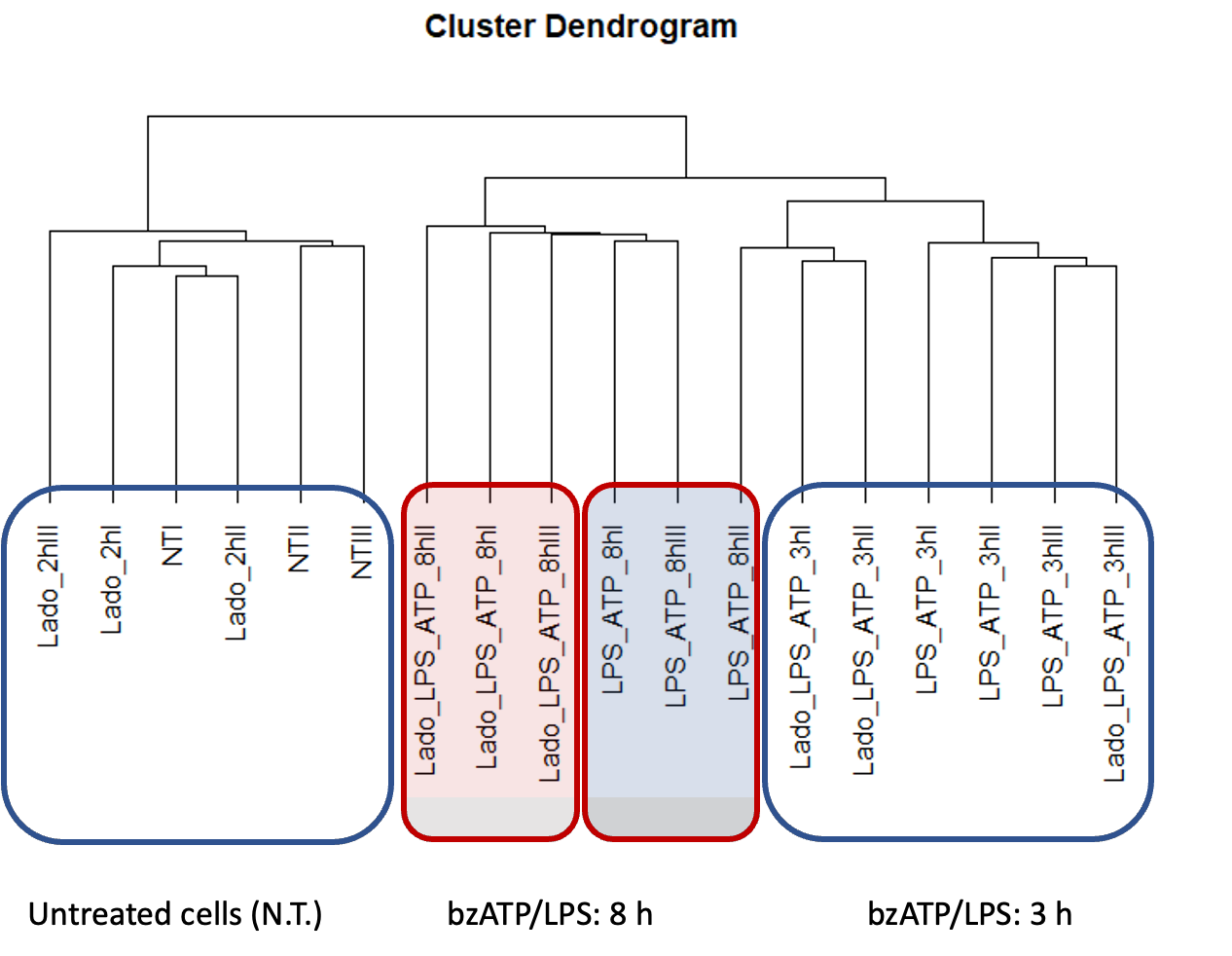

### ***Figure S1. Dendogram of the 18 tests and their control experiments. The dendogram used all mapped miRNAs (372). Each experiment is marked as I, II and III. The incubation of ladostigil (Lado) cased no effect as shown for N.T.) and similarly, the effect of shows that ladostigil in 3 h is negligible.***

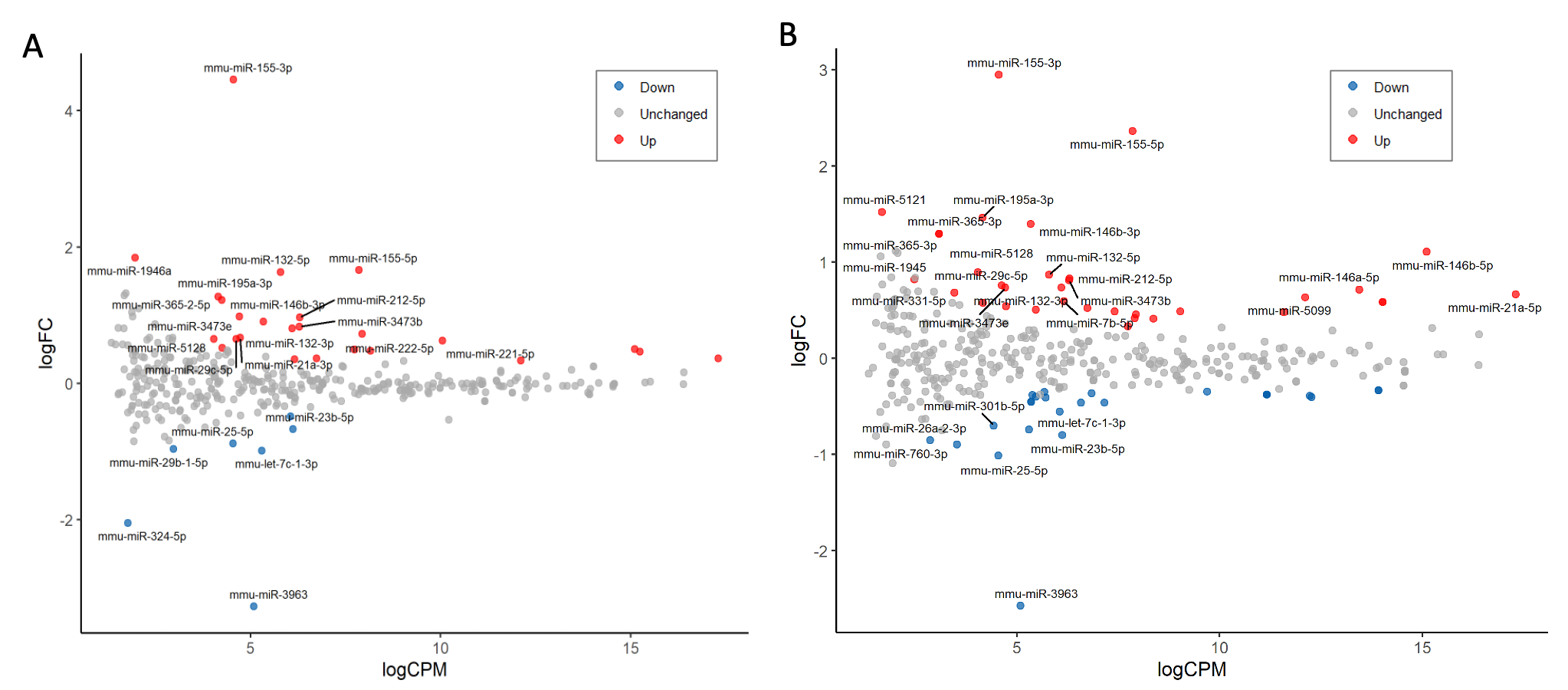

**Figure S2.** *The effect of microglial activation.* ***(A)*** *MA plots of the log2(FC) relative to the CPM (in log) for the 372 identified miRNAs in an activated setting (bzATP/LPS) 3 h post activation.* ***(B)*** *MA plots of the log2(FC) relative to the CPM (in log) miRNAs in an activated setting (bzATP/LPS) 8 h post activation relative to non-treated cells (N.T.).*
